# A Comprehensive Mathematical Model of Surface Electromyography and Force Generation

**DOI:** 10.1101/273458

**Authors:** Eike Petersen, Philipp Rostalski

## Abstract

The purpose of this article is to provide a unified description of a comprehensive mathematical model of surface electromyographic (EMG) measurements and the corresponding force signal in skeletal muscles. The model comprises motor unit pool organization, recruitment and rate coding, intracellular action potential generation and the resulting EMG measurements, as well as the generated muscular force during voluntary isometric contractions. It consolidates and extends the results of several previous publications that proposed mathematical models for the individual model components. A parameterization of the electrical and mechanical components of the model is proposed that ensures a physiologically meaningful EMG-force relation in the simulated signals. Moreover, a novel nonlinear transformation of the excitation model input is proposed, which ensures that the model force output equals the desired target force. Finally, an alternative analytical formulation of the EMG model is proposed, which renders the physiological meaning of the model more clear and facilitates a mathematical proof that muscle fibers in this model at no point in time represent a net current source or sink.

Neuromuscular physiology is a vibrant research field that has recently seen exciting advances. Many previous publications have focused on thorough analyses of particular aspects of neuromuscular physiology, yet an integration of the various novel findings into a single, comprehensive model is missing. A consistent description of a complete physiological model as presented here, including thorough justification of model component choices, will facilitate the use of these advanced models in future research. Results of a numerical simulation highlight the model’s capability to reproduce many physiological effects observed in experimental measurements, and to produce realistic synthetic data that are useful for the validation of signal processing algorithms. The model is based on recent advances in the understanding of muscular physiology and hence also applicable for analyzing the influence of various physiological and measurement setup parameters on the measured force and EMG signals.

## 1 Introduction

Electromyography (EMG) denotes the measurement of the electrical fields generated by the electrophysiological processes that lead to muscle fiber contraction. EMG is highly relevant for a number of clinical and scientific application fields, since it enables monitoring and analysis of a muscle’s electromechanical properties and state, both of which would otherwise remain mostly inaccessible. Surface electromyography (sEMG) denotes the noninvasive measurement of electrical muscle activity by means of electrodes placed on the skin surface, as opposed to the traditional measuring method using needle electrodes. For more background information on sEMG, its analysis and many of its applications, refer to, e.g., Merletti and Parker (30); Merletti and Farina (31).

Mathematical models of sEMG are highly useful, on the one hand to advance understanding of the underlying physiological processes, and on the other hand to analyze the sensitivity of sEMG measurements to various physiological and technical parameters, and to test and validate sEMG signal processing algorithms. Over the past decades, researchers have pursued a number of different approaches for the modelling and simulation of different aspects of sEMG measurements. Phenomenological (23; 28; 29; 50) as well as physiologically motivated (13–15; 17; 19; 20; 34) models have been proposed and analyzed. Overviews can be found in McGill (29); Rodriguez-Falces et al. (47); Stegeman et al. (51) and Merletti and Farina (31).

Particular emphasis has been placed on modelling the electric signal produced by a single contraction of a single muscle fiber, the so-called single fiber action potential (SFAP). Classically, simplified dipole, tripole or quadrupole models have been employed for modelling the propagation of the action potential along a contracting muscle fiber (21; 30; 32; 41). A more general model has been proposed by Dimitrov and Dimitrova (15), and this model has been successfully employed, modified and combined with various other models for the remaining physiological processes in a number of publications (14; 17; 19; 55).

In the present article, the SFAP model originally proposed by Dimitrov and Dimitrova (15), and subsequently extended by Farina and Merletti (17), is combined with the well-known motor unit (MU) pool organization model of Fuglevand et al. (20) and the twitch force parameterization used by Raikova and Aladjov (43). Recent results regarding the modelling of MU rate coding and recruitment (10) and the variability of the inter-spike intervals (35) are incorporated. Care is taken in particular to achieve a consistent parameterization of the electrical and the mechanical components of the model, resulting in a realistic EMG-force relationship of the simulated muscle. Moreover, a novel nonlinear transformation of the rate coding model is proposed, which ensures that the desired output force is matched by the model. Finally, an alternative analytical formulation of the SFAP model of Farina and Merletti (17) is proposed, which renders the physiological meaning of the model more clear. Based on this alternative formulation, a proof is provided that in this model no fiber represents a net current source or sink at any point in time, which is a physiologically plausible property due to the quasi-static behavior of action potential generation (41). It is also in accordance with the predictions of the reknown Hodgkin-Huxley model (27).

In the following section 2, all components of the mathematical model are presented briefly, yet completely, and in a unified way. Several mathematical properties of the model are derived, and the mentioned alternative formulation of the model of Farina and Merletti (17) is proposed. Results of a numerical simulation of the model are presented in section 3 and are assessed with respect to their physiological plausibility. Finally, section 4 concludes the article with a brief summary. A preliminary version of some of the results regarding the sEMG model presented in this article has previously been the subject of a conference publication (37).

## 2 Mathematical model

The fundamental functional unit of a skeletal muscle is the motor unit (MU), comprising a motor neuron and all muscle fibers innervated by that neuron. The following sections introduce mathematical models of the electrical and mechanical properties of MUs, as well as their organization in a muscle. Figure 1 shows a graphical summary of the main model components and their interactions and may provide a useful reference for the reader while following along the description of the model.

**Figure 1:**
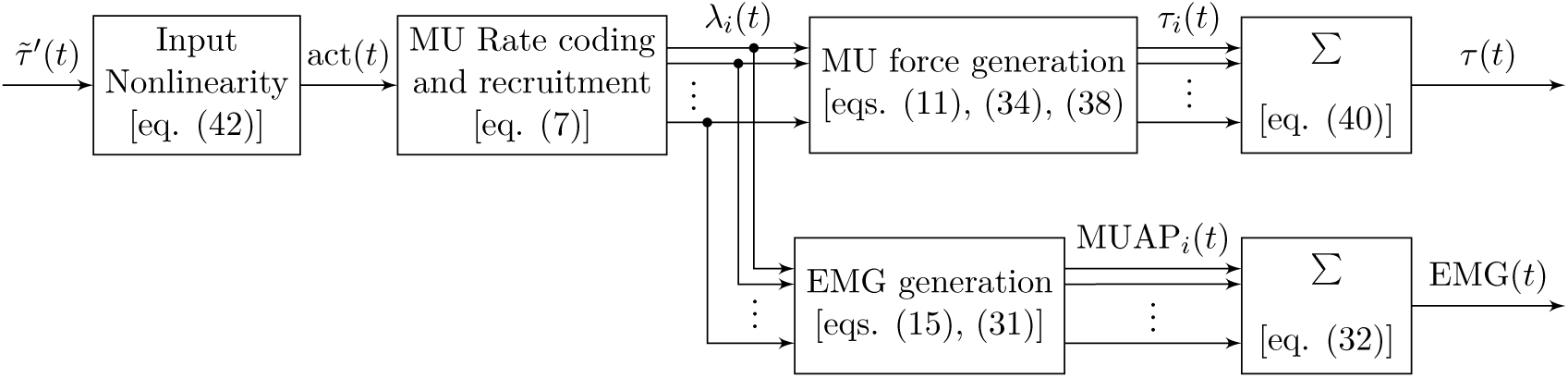
Block diagram illustrating the main components of the proposed model of muscular force generation. The model input is the normalized desired muscle force 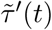 with values in [0, 1], the model outputs are the total generated muscle force *τ* (*t*) and the measured EMG signal EMG(*t*). Firing rates of individual MUs are denoted by *λ*_*i*_(*t*), the current force contribution of each MU by *τ*_*i*_(*t*), and the current EMG contribution of each MU by MUAP_*i*_(*t*).

### 2.1 Motor unit pool structure

Every muscle consists of a number *n* of MUs. Each MU has various mechanical and electrical properties, most of which have been found to be closely related by means of the *size principle* (22): MU size as measured by the number of fibers contained in the MU is roughly proportional to force twitch amplitude, EMG twitch amplitude, and recruitment threshold. This means that larger MUs are only activated at higher levels of desired muscle force compared to smaller units, but they also add larger force and EMG contributions to the muscle output once activated. Recent results have shown that the electrical twitch conduction velocity *ν* is linearly related to the recruitment threshold, and hence can be considered a size principle parameter aswell (11). Setting one of these parameters to a particular value for a given MU also determines the remaining parameters.

The recruitment thresholds appear to follow a continuous distribution with many MUs attaining a small recruitment threshold, and few large MUs only being recruited at high activation levels (44). This behavior is captured well by the exponential model proposed by Fuglevand et al. (20), which assigns the recruitment thresholds

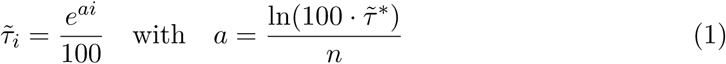

to MUs *i* = 1, *…, n*, where 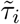 denotes the minimum fraction of total muscle force at which the MU is recruited, and 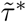 denotes the point of full recruitment, i.e., the relative level of total muscle force at which all MUs are recruited. Alternatively, following the formulation of De Luca and Contessa (8), the thresholds can also be modelled as

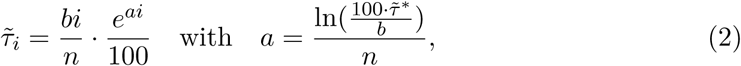

where *b* denotes a scaling factor that influences the shape of the distribution. The latter model results in a more gradual slope compared to the first one and has been used by De Luca and Contessa (8) for modelling the characteristics of the Vastus Lateralis (VL) muscle (with *b* = 20). The choice of one of equations (1) and (2) in general should be based on the characteristics of the specific muscle under consideration.

With the recruitment thresholds set, peak twitch forces are calculated as a linear function of the recruitment thresholds following

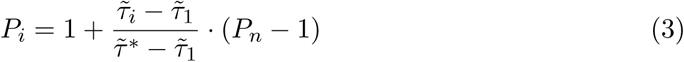

as proposed by Contessa and De Luca (7), where the twitch peak range (*P*_*n*_*/P*_1_) is typically large, e.g., *P*_*n*_*/P*_1_ = 130 for the First Dorsal Interosseus (FDI) muscle (7). Equivalently, the number *η* of innervated muscle fibers, which appears to be the main factor influencing MU twitch force (20), can be modelled directly proportional to the peak twitch force and hence also to the recruitment threshold, i.e.,

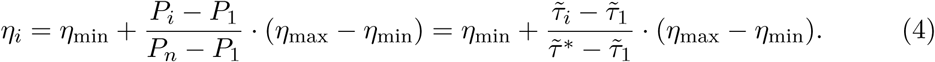

The same is true for the electrical twitch conduction velocity *ν*, as noted above (11). Note that equation (4) determines the relation between the electrical and mechanical properties of a MU, as the number of fibers in a MU determines the amplitude of the electrical twitch response in this model. In equation (4), a linear relationship between the two has been assumed. However, it has been shown that in some muscles, the electrical twitch amplitude may rather be related to the square root of the force twitch amplitude (57); such a relation can easily be implemented into the model by modifying equation (4) to

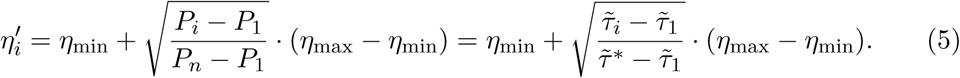

### 2.2 Geometrical distribution of motor units and muscle fibers

Muscle fibers belonging to the same motor unit are distributed in a territory that may span a large portion of the muscle cross-section (3; 4; 53). The territories of the different MUs overlap, leading to fibers belonging to multiple MUs being intermingled (3; 53). Motor unit territories have been found to attain an irregular round shape (3), whence we propose the use of an elliptic model for the MU cross-sections. With the elliptic axis ratio being fixed, the MU cross-sectional area—and thus the axes lengths—is calculated by dividing the number of innervated fibres *η* by the desired MU fiber density *ρ* ibers*/*area):

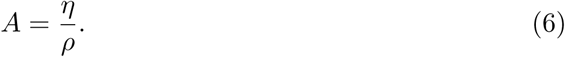

The midpoints of all MUs are then distributed uniformly over the muscle cross-section. Note that without further assumptions, the above model directly leads to overlapping regions between MUs, which is a desirable feature, as noted above. Finally, fibers belonging to the MU are then again distributed uniformly inside the elliptic MU crosssection. The model is equivalent to the propositions of Fuglevand et al. (20), except for the division of the muscle into multiple parts to reduce fiber density variability, and the fact that they used circular MU territory shapes as opposed to the more flexible elliptic shape proposed here.

A key decision when modelling random MU placement concerns the treatment of muscle boundaries. Those parts of MU territories that exceed the muscle territory must be cut off and the question remains how to account for this loss in MU territory. Several approaches to solving the problem are conceivable (47):

1. All fibers belonging to the MU are placed in the remaining parts of the MU territory. This approach leads to an increase in the fiber density of boundary MUs, and hence also in the overall fiber density towards the muscle boundaries.
2. The number of fibers innervated by the MU is reduced proportionally. This approach keeps the assigned fiber density constant but reduces the number of innervated fibers and thus the size of boundary MUs.
3. The axes lengths of the elliptic MU region are adjusted in such a way as to keep the MU area at the desired value in spite of the cut-off. This keeps the number of fibers and the fiber density at the desired values but likely leads to strongly increased overall fiber density towards the muscle centre, due to many adjusted MU regions overlapping there.

There are advantages and disadvantages to each approach, and it does not yet seem to be clear if one of the proposed approaches is generally superior to the others, or which approximates reality best (47). However, muscle fiber diameters appear to be approximately constant throughout a muscle (24; 49), whence a constant fiber density throughout the muscle cross-section seems desirable. To this end, the second of the above approaches has been pursued here. In order to avoid a high variability of the fiber density due to the random MU placement, it is advisable to divide the muscle cross-section into *M* parts of equal size, and then distribute *n/M* MUs uniformly in each part.

Reducing only the number of fibers in MUs close to the muscle boundary without adjusting the other MU parameters as well would disturb the relation between the electrical and mechanical properties of these MUs. To account for this disturbance, the recruitment threshold 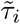, the peak twitch force *P*_*i*_ and the electrical twitch conduction velocity *ν* have been recomputed following equations (3) and (4) for these MUs. Note that this, in turn, distorts the exponential distribution of MU parameters described by equations (1) and (2). This, however, was considered less grave than a disturbed electromechanical relationship in a significant number of MUs.

An alternative model of MU placement has been proposed by Navallas et al. (36). They explicitly considered the optimization problem of minimizing the variability of muscle fiber density throughout the muscle, regardless of variances in MU fiber density and while maintaining the exponential relationship (1). Their model has recently been extended to account for regionalized MU placement (46). While all of these are desirable properties of a MU placement algorithm, the variability of the fiber density may also be reduced by the much simpler division of the muscle region into distinct parts and random placement proposed above; MU regionalization could be implemented using this same division into distinct parts; and the distortion to the exponential relationship (1) when using this algorithm has been found to be rather small in practice, see section 3. In summary, both the placement algorithms of Navallas et al. (36) and the one proposed here represent viable modelling choices, with the former probably being preferrable if the particular influence of different aspects of MU geometry is of interest, and the latter being a much simpler algorithm.

### 2.3 Firing rates

Following the experimental results of Milner-Brown et al. (33), firing rates have traditionally been modelled as a linear function of excitatory drive (16; 20). This simplified model has recently been superseeded by the linear-exponential firing rate model proposed by De Luca and Hostage (10). In this model, the firing rate of a MU is described as

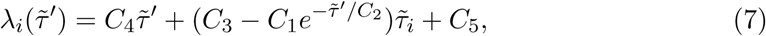

where 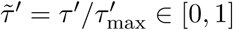 denotes the relative level of total desired muscle force *τ* ′, and the remaining constants are shape parameters. This model reproduces many phenomena observed experimentally, such as the *onion-skin phenomenon*: MUs recruited first appear to attain a higher firing rate throughout the whole contraction than those MUs recruited later (9). Interestingly, however, the model also misses an essential feature of the model proposed earlier by Erim et al. (16), namely, the (experimentally observed (9)) increased slope of the firing rate characteristics at excitation levels exceeding the point 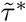 of full recruitment. We will come back to this issue in section 2.8. For example values of the shape parameters for different muscles, refer to De Luca and Hostage (10).

Note that although the model is defined as a function of the desired total muscle force, it does not guarantee in any way that this force level is actually attained by the muscle. This inconsistency will be addressed in section 2.8 of this article by introducing a suitably defined nonlinear input transformation act 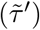 that is then used as the input to the rate coding model presented above, instead of using the desired muscle force level 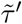 directly. One benefit of the nonlinear input transformation proposed in section 2.8 is that it effectively results in an increased slope of the firing rate characteristics at excitation levels exceeding 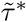, as has been observed experimentally (9).

### 2.4 Firing instants

Given the instant *t*_*i*(*j*−1)_ of the last firing of MU *i* and the time course of the MU’s firing rate *λ*_*i*_, the next firing instant *t*_*ij*_ can be calculated. To model the stochastic distribution of the inter-spike intervals, these are assumed to follow a normal distribution with a coefficient of variation that decreases with increasing activation, following

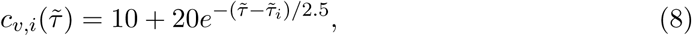

as proposed by Moritz et al. (35). The *j*^th^ inter-spike interval ISI_*j*_ then is drawn from

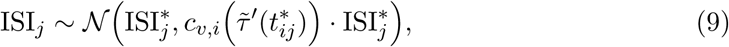

where the mean 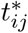 of the time of the next firing event and the corresponding mean inter-spike interval 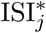 are obtained by solving

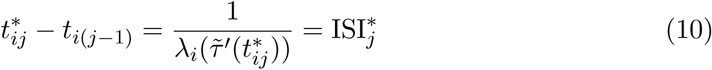

for 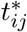 as proposed by Fuglevand et al. (20), with 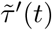 denoting the normalized requested muscle force level at time *t*. The time *t*_*ij*_ of the *j*^th^ firing event is then calculated as

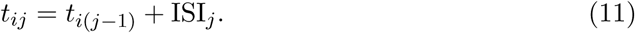

Note that several distributions other than the normal distribution have been proposed for modelling the distribution of the inter-spike intervals (1; 25). For reasons of simplicity, and as the influence of the shape of the distribution of the inter-spike intervals on the overall EMG and force signals found by Barry et al. (1) appeared to be rather negligible, a normal distribution is used in the simulation described in section 3.

### 2.5 Intracellular action potential propagation

The propagation of an intracellular action potential (IAP) from the neuromuscular junction (NMJ) of a muscle fiber along both directions towards the two fiber ends can be modelled by representing the actively firing fiber by a distributed current source and sink. In the model originally proposed by Dimitrov and Dimitrova (15), this distributed fiber membrane current source *î*(*z, t*) is composed of two propagating wave fronts and localized contributions at the NMJ and the two fiber ends. These localized contributions model the IAP generation and extinction process. In the formulation of Farina and Merletti (17), the model reads

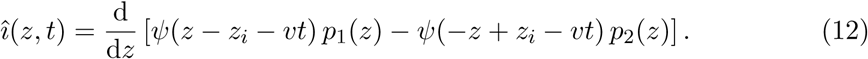

Here, *z* denotes the spatial variable along the muscle fiber, *z*_*i*_ the location of the NMJ, *L*_1_ and *L*_2_ are the distances between the innervation zone and the right and left tendon, respectively, and *ν* denotes the IAP’s propagation velocity. Moreover,

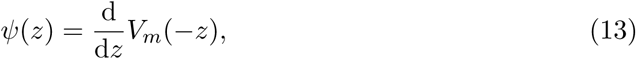

denotes the voltage gradient across the fiber membrane along the fiber axis, where the function *V*_*m*_(*z*) prescribes a model for the trans-fiber membrane voltage wave shape and can be chosen arbitrarily to match simulated or measured data. Refer to Plonsey and Barr (41) for details on the significance of *V*_*m*_(*z*). Here, the analytical model function

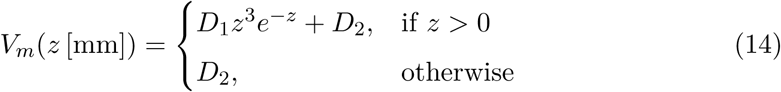

with *D*_1_ = 96 mV mm^−3^ and *D*_2_ = −90 mV will be used, as originally proposed by Rosenfalck (48) and as has been done by Farina and Merletti (17).

The IAP model in equation (12) can be shown to be equivalent to choosing

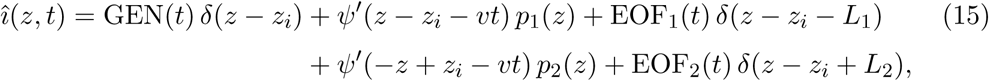

with the Dirac distribution *δ*, the end-of-fiber components

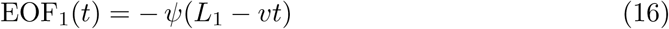

and

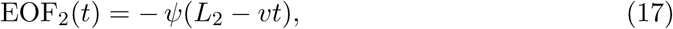

and the potential generation component

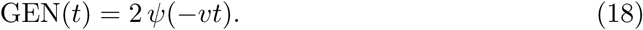

Here, again, the end-of-fiber components describe the IAP extinction process at the fiber ends, and the potential generation component models the influence of IAP generation at the innervation zone on the membrane current.

This formulation renders – to the authors’ opinion – the structure of the model more obvious, by clearly distinguishing between propagating and non-propagating signal components, and by revealing the non-smoothness of the resulting distributed current source, the latter following from the presence of the stationary Dirac distributions at the two fiber ends and the location of the innervation zone. The equivalence of the two formulations of the model is summarized in the following lemma, the proof of which is given in the appendix.

#### Lemma 1.

*The expressions given in equations* (12) *and* (15) *to* (18) *are equivalent, assuming that ψ ∈ 𝒞* ^*∞*^.^*1*^

Interestingly, one can also show that this IAP model ensures

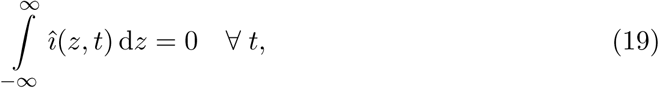

which implies that the fiber does not represent a net current source or sink at any point in time. This property is well motivated by physiology, considering that the invoked electrodynamical processes can be considered quasi-static (41), and it is also in accordance with the predictions of the reknown Hodgkin-Huxley model for action potential propagation (27). It is precisely a result of the presence of the three Dirac distributions in the model equation (15), as these have the combined effect of collecting all remaining currents exerted by the intermediate fiber sections due to their higher potential. This result is the subject of the following lemma, the proof of which again is deferred to the appendix.

#### Lemma 2.

*For compactly supported ψ*(*z*), *the IAP model given in equations* (15) *to* (18) *yields a formulation of î*(*z, t*) *that satisfies equation* (19).^*2*^

### 2.6 EMG measurements

Biological tissues can be considered volume conductors (41). The existence of an electric field implies the existence of electric currents travelling through the tissue, and vice versa.^3^ Due to the comparably low rate of change of physiological systems, it is justified (41) to assume these time-varying electric fields to behave as if they were static at each instant of time, whence they are called *quasi-static*. This assumption amounts to a neglection of the capacitive properties of the tissues. Accordingly, as for static fields, the electric field in a physiological volume conductor is considered equal to the negative gradient of a scalar potential *φ*, namely,

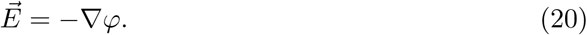

By Ohm’s law, the current density (current per unit of cross-sectional area) in a volume conductor is proportional to the electric field, that is,

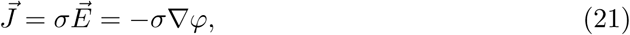

where *σ* denotes the conductivity of the medium. Defining a distributed current density source *I* throughout the region of interest, the divergence of the current density is constrained by

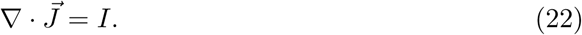

Combining equations (21) and (22) and assuming a homogeneous, isotropic medium yields Poisson’s equation for the diffusion of the potential, namely,

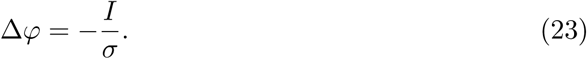

In the following, the electric field generated by point sources in planar tissue layers will be considered as a model for flat and large muscles, such as the *recti abdominis* simulated in section 3. The muscle layer is assumed to be infinitely extended and planar, and to be covered by an infinitely extended planar layer of fat and an infinitely extended planar layer of skin. Muscle tissue is considered anisotropic in order to reflect the difference in conductivity between currents along the muscle fiber axis and currents across the muscle fiber axis, whereas fat and skin tissue are considered isotropic. Muscle fibers are assumed to run along the *z* direction, with the *x* and *z* dimensions spanning the skin plane, and the *y* dimension being orthogonal to the skin plane, positive vectors pointing outwards.

The geometrical set-up described above has been analyzed by Farina and Rainoldi (18). For a point source of strength *Î* located at (0, *y*_0_, 0), the authors derive the 2-D spatial Fourier transform of the resulting potential distribution at the skin surface to be

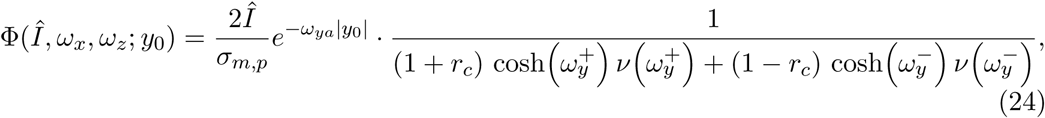

with the abbreviations

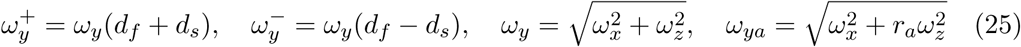

and

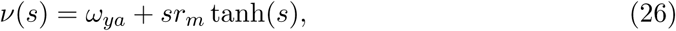

where *ω*_*x*_ = 2*πf*_*x*_ and *ω*_*z*_ = 2*πf*_*z*_ denote the spatial angular frequencies in the *x* and *z* directions, respectively. The coefficients

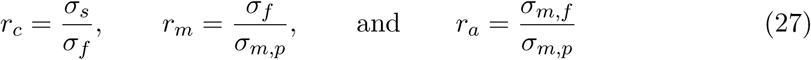

specify ratios of the different tissue conductivities. Finally, *y*_0_ denotes the depth of the point source in the muscular tissue, *d*_*f*_ the thickness of the fat layer and *d*_*s*_ the thickness of the skin layer.

Equation (24) directly yields an analytic description of the 2-D spatial transfer function of the volume conductor *via*

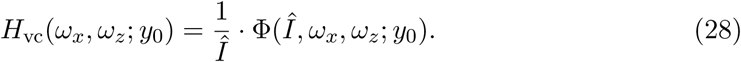

EMG measurements are usually taken differentially between a set of electrodes. Consider a regular grid of *R×S* electrodes with interelectrode distances *d*_*x*_ and *d*_*z*_, espectively, where *R* = *R*_*a*_ + *R*_*b*_ + 1 and *S* = *S*_*a*_ + *S*_*b*_ + 1. The variables subscripted by *a* and *b* denote the number of electrodes on the two sides of an arbitrarily chosen reference electrode. The grid is assumed to be aligned parallel to the *z* axis. Assigning weights *ζ*_*k𝓁*_ to the electrodes and assuming all electrodes to attain the same transfer function, the (spatial) transfer function from a given surface potential distribution to the potential measured by such an electrode configuration at each point on the surface is given by (17)

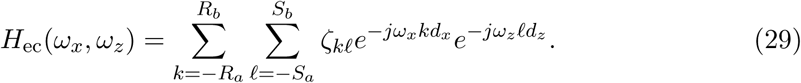

For the transfer function of a single electrode, arbitrary model assumptions can be made. For details, refer to, e.g., Merletti and Parker (30).

Concatenating the spatial transfer functions *H*_vc_ of the volume conductor, *H*_ec_ of the electrode configuration and *H*_ele_ of the electrodes themselves, the global transfer function of the combined system emerges as

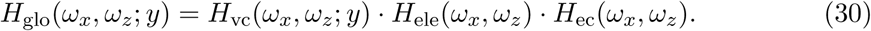

From this, the 2-D potential distribution on the skin surface can generally be calculated as

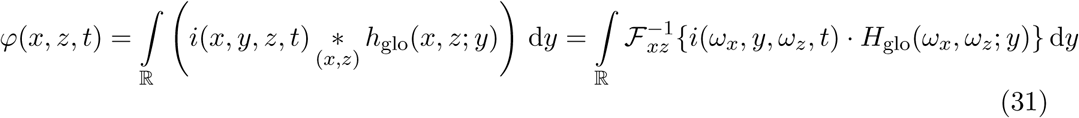

where *i*(*ω*_*x*_, *y, ω*_*z*_, *t*) = *ℱ*_*xz*_{*i*(*x, y, z, t*)} is the 2-D Fourier transform of the current density source *i*(*x, y, z, t*), and *\**_(*x,z*)_ denotes 2-dimensional convolution in the *x* and *z* variables. For a particular electrode location on the skin surface and a muscle fiber following a straight line parallel to the skin surface, equation (31) simplifies, and the resulting single-fiber action potential (SFAP) *φ*(*t*) can be calculated numerically (17; 38). One can prove that in this case the integration kernel only has removable singularities, which ensures the convergence of a numerical integration scheme (38). Figure 2 shows exemplary SFAPs resulting from the evaluation of equation (31) for such fibers using nested numerical integration schemes. Note that while the above derivation has been performed for the case of planar volume conductors, similar models have been derived for cylindrical volume conductors – much more appropriate for the simulation of limb muscles – as well (19).

**Figure 2:**
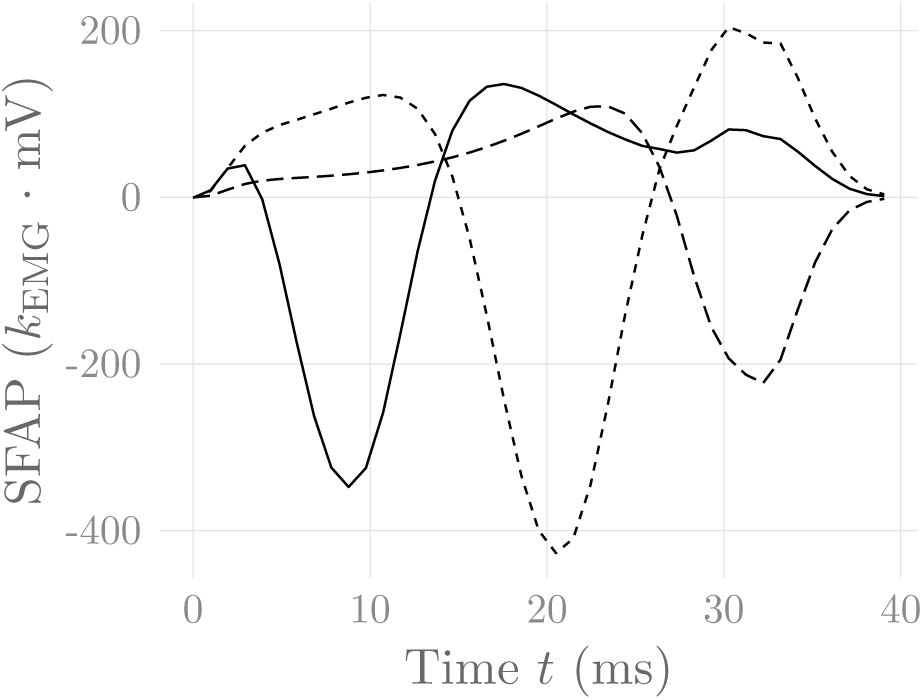
Simulated SFAPs evoked by a single firing muscle fiber as detected by three surface electrodes positioned close to the NMJ (solid), in between the NMJ and the fiber end (short dashes) and above the fiber end (long dashes).

The numerical solution of the (simplified version of) equation (31) is computationally moderately expensive. Fortunately, this only has to be done once for each fiber before the actual simulation, in order to calculate the SFAPs of all fibers. During the simulation, multiple shifted versions of these SFAPs are then superposed to generate the actual EMG measurement

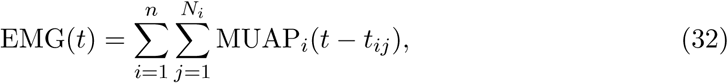

where *n* denotes the number of MUs, *N*_*i*_ the number of firing events of that MU, and

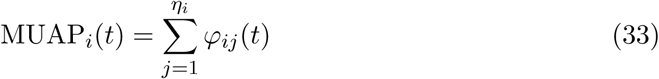

the motor unit action potential, which is obtained by summing over the contributions, i.e., the SFAPs, of all muscle fibers belonging to MU *i*. Figure 2 shows exemplary SFAPs simulated using a numerical implementation of the above model. The simulated SFAPs correctly reproduce the dependency of the SFAP shape on the relative position of the recording electrode and the depth of the muscle fiber, as well as the distinction between propagating and localized signal components at the NMJ and the two fiber ends. In particular, the experimentally observed end-of-fiber extinction signals are included.

### 2.7 Force twitches

Each MUAP generates a corresponding force contribution, denoted as a force twitch *f*_*i*_(*t*). Most previously proposed muscle models (13; 20) have employed the force twitch parameterization of Milner-Brown et al. (33), where a force twitch is completely described by its twitch rise time *T*_ri_ and its peak twitch force *P* This model, however, does not allow setting the half relaxation time *T*_hr_ independently of *T*_ri_, which is essential for modelling different muscle fiber types (refer to Fig. 1.2 in (30), for example). For this reason, we propose the use of a different model. Raikova and Aladjov (43) used a model of increased expressivity, which is obtained by introducing an additional degree of freedom, and which results in the following force twitch model:

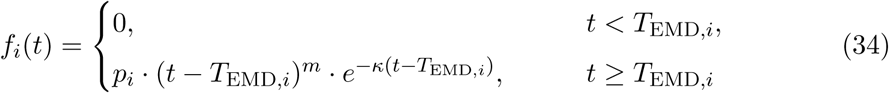

with

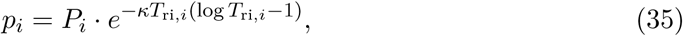

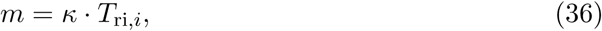

and

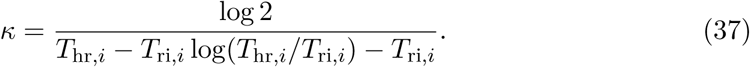

This model satisfies *f*_*i*_(*T*_ri,*i*_) = *P*_*i*_, and *f*_*i*_(*T*_hr,*i*_) = *P*_*i*_*/*2. The parameter *T*_EMD_ denotes the electromechanical delay between the onset of electrical and mechanical activity of the motor unit. The three parameters *T*_ri_, *T*_hr_ and *T*_EMD_ are sampled from a Weibull distribution for each MU, as proposed (for the former two) by Contessa and De Luca (7). Figure 3 shows some exemplary force twitches generated by this model. Note that fixing *m* = 1 and freely selecting *p*_*i*_ and *κ* results in the force model of Milner-Brown et al. (33).

**Figure 3:**
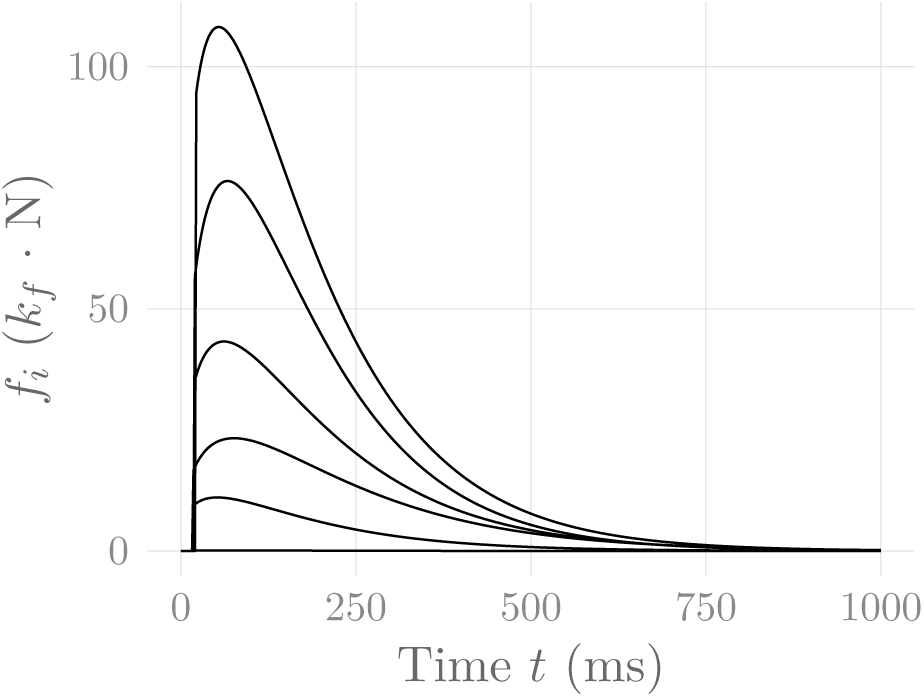
Force twitches generated by every tenth MU of the muscle simulated in section 3. These are the same MUs for which the rate coding characteristics are shown in fig. 4.

Kernell et al. (26) have found experimentally that there is a nonlinear relationship between generated isometric muscle force and firing rates of MUs: At high firing rates, the force twitch amplitude decreases. This nonlinear relationship is included in the proposed model by scaling individual force twitches in the impulse train of a particular MU by a factor

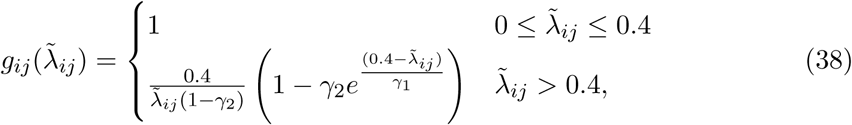

with *γ*_2_ and *γ*_1_ constant muscle parameters, as proposed by Contessa and De Luca (7). Here, *g*_*ij*_ denotes the gain factor assigned to the *j*th firing of MU *i*, and 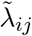 is the normalized instantaneous firing rate at that firing event:

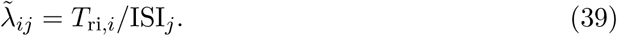

Finally, the total force generated by a muscle is calculated as the superposition

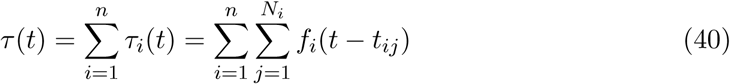

of the individual force twitches of all MUs, with *n* the number of MUs in the muscle, *τ*_*i*_(*t*) the force contribution of MU *i* over time, *N*_*i*_ the number of firing events of MU *i*, and *t*_*ij*_ the *j*^th^ firing instant of MU *i*, calculated following equation (11).

### 2.8 Excitation-force relationship

Figure 1 illustrates the model of muscular force generation described so far, introducing a muscle activation signal act ∈ [0, 1]. Motor Unit firing rate models such as the model of De Luca and Hostage (10) described in section 2.3 usually define the firing rate as a function of the desired normalized muscle force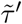, i.e., they choose act 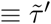. This is a consequence of the fact that these models are derived from experimental measurements of MU firing rates at different muscle force levels. Now, defining the firing rates *λ*_*i*_ of all MUs (e.g. as described in section 2.3) and also defining the force generating properties of these MUs (e.g. as described in section 2.7) uniquely determines the generated muscle force *τ*, see fig. 1. It is, however, by no means guaranteed that the generated normalized muscle force 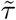 will be equal to the force level 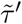 that has been used as an input to the firing rate model. In other words, the generated muscle force output does not match the desired muscle force.

Dideriksen et al. (14); Venugopal et al. (54) dealt with this problem by introducing a simple PID controller for adjusting the model input, such that the error between desired and actual force output is minimized. This corresponds to the introduction of a feedback loop in fig. 1. In their model, parameters change over time (simulating fatigue), rendering the introduction of such a feedback loop an elegant solution to the problem of time-varying input-output consistency. Contessa and De Luca (7) introduced a similar force feedback loop, although they implemented the feedback in an offline instead of online fashion, updating the input signal in hindsight and re-running the simulation if the force output deviated too strongly from the desired output. While all three articles (7; 14; 54) include some remarks on physiological feedback processes, neither model was meant to replicate properties of actual physiological feedback control, but rather to account for the input-output inconsistency of the respective model. Although there certainly is some feedback element in physiological force control, it would appear reasonable that this feedback is mainly necessary to deal with external disturbances and changing muscle properties, not to account for a static input-output inconsistency. For these reasons, we propose a novel solution to this problem which does not require the introduction of a feedback loop. Our approach is based on a static input nonlinearity.

Denoting the desired normalized muscle force by 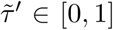, our desire is to choose act 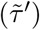 such that

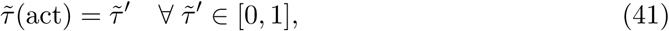

where 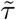(act) is the normalized generated force output. This can be achieved by calculating 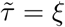(act), i.e., the normalized generated muscle force as a (nonlinear) function *ξ* of the activation input. Utilizing

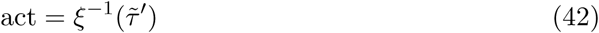

as the activation input to the firing rate model then yields a simulation model that satisfies condition (41). Averaging over individual firing events, the mean generated muscle force is given by

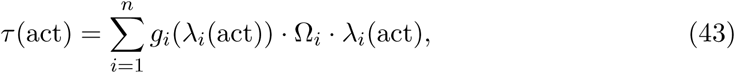

where

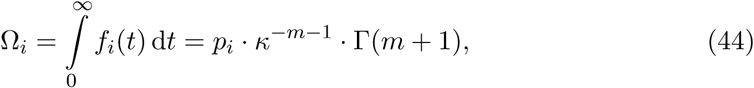

with Γ(*x*) the Gamma function, denotes the total impulse generated by a single force twitch *f*_*i*_(*t*) of MU *i*, and *g*_*i*_ denotes the nonlinear force gain factor defined in equation (38). By evaluating

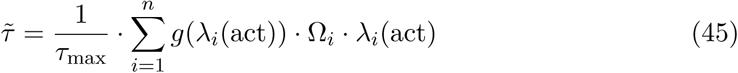

for different values of the activation level act, one can determine 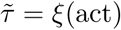. Employing 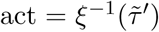 as the activation input to the firing rate model then yields a simulation model that satisfies

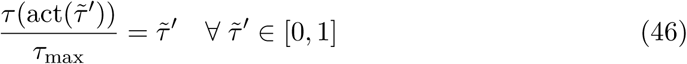

in the mean over time. Condition (46) ensures that while there may be differences between desired and generated muscle force at individual time instants due to the stochastic nature of the firing instants (see section 2.4), the two forces agree on average. Finally, note that the introduction of a static input nonlinearity leads to a distortion of the firing rate model, which now no longer receives 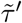 as an input, but rather its nonlinear transformation 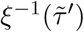. The influence of this distortion will be exemplified in the following section, and its plausibility will be discussed.

## 3 Simulation results

Using the mathematical model presented in the previous section, a numerical simulation of the rectus abdominis muscle has been implemented as a test scenario, using the R programming language (42). In total, 300 MUs have been simulated, each consisting of between 30 and 150 muscle fibers, organized into three separate muscle bellies of each of the two recti (left and right). Single differential detection was assumed for the EMG signals, with the two electrodes placed on the linea alba between the second and third belly. Where available, parameter values available in the literature have been chosen (12; 45; 52). A series of constant-force isometric contractions at progressively increasing force levels has been simulated, each lasting for 4 s. In order to assess the effect of the the inherent randomness of the MU placement algorithm, the muscle generation procedure and the subsequent simulation each have been executed seven times. The simulated signals are available at the Dryad data repository (39).

Figure 4 shows a comparison of the rate coding model with and without the static input nonlinearity introduced in section 2.8. It can be observed that the input nonlinearity introduces an increase in the slope of the firing rate curves of all MUs starting at the point of full recruitment. This is in perfect agreement with the experimental findings of Erim et al. (16) and also makes sense from an intuitive point of view: Once all MUs are recruited, firing rates must increase faster than before in order to achieve an increase in muscle force output. It hence appears that the introduction of this static input nonlinearity, calculated by considering all model parameters together, increases the degree of similarity between experimental observations and the rate coding and recruitment model of De Luca and Hostage (10). The non-smoothness of the adjusted rate coding characteristics is a consequence of the same non-smoothness in the original activation-force relationship, due to new MUs being recruited at discrete levels of excitation. Accordingly, the non-smoothness of the adjusted rate coding characteristics is required to obtain a smooth input-output force relationship.

**Figure 4:**
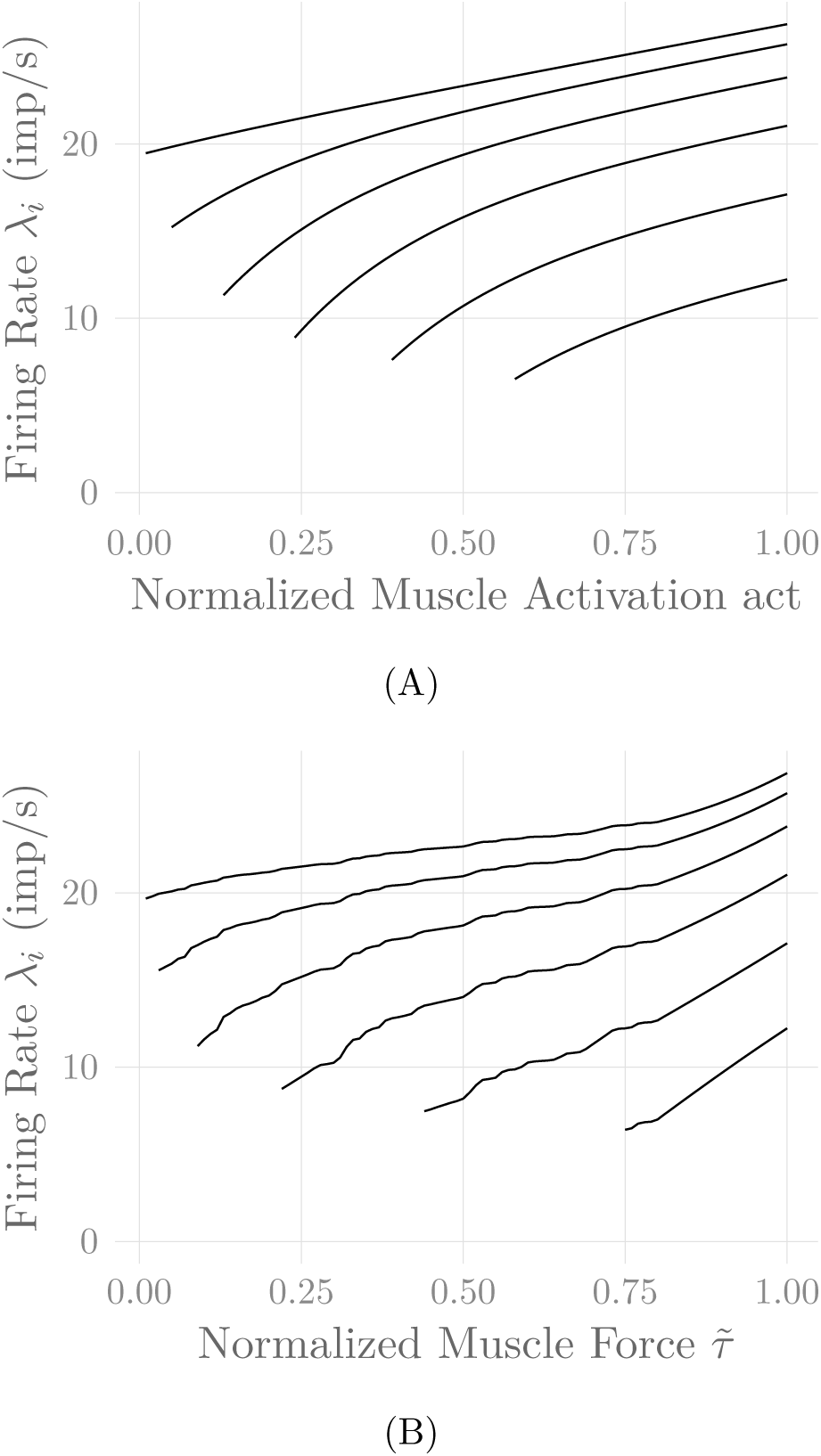
Firing rates of every tenth MU in one belly of one of the simulated *recti*, (A) as a function of the rate coding model input act, and (B) as a function of the desired output muscle force 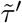. The point of full recruitment was set to 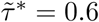. Note that the application of the nonlinear input transformation 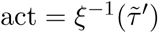 has shifted the point of full recruitment to a higher force level.

Figure 5 shows the amplitude distribution of the simulated EMG signal, which resembles a smoothed Laplacian distribution. This type of distribution has been reported previously for the amplitude distributions of real measurement signals (5). Figure 6 displays the simulated series of isometric contractions; fig. 7 shows the coefficient of variation and the standard deviation of the simulated force signal at different levels of muscle activation. Both force variability graphs nicely resemble those reported by (1) for experimental measurements of index finger force steadiness. Finally, fig. 8 shows the sEMG-Force relationship calculated over the simulated series of isometric contractions, both for the linear electromechanical relationship described by equation (4) as well as for the square root relationship described by equation (5). Both graphs behave as expected (57).

**Figure 5:**
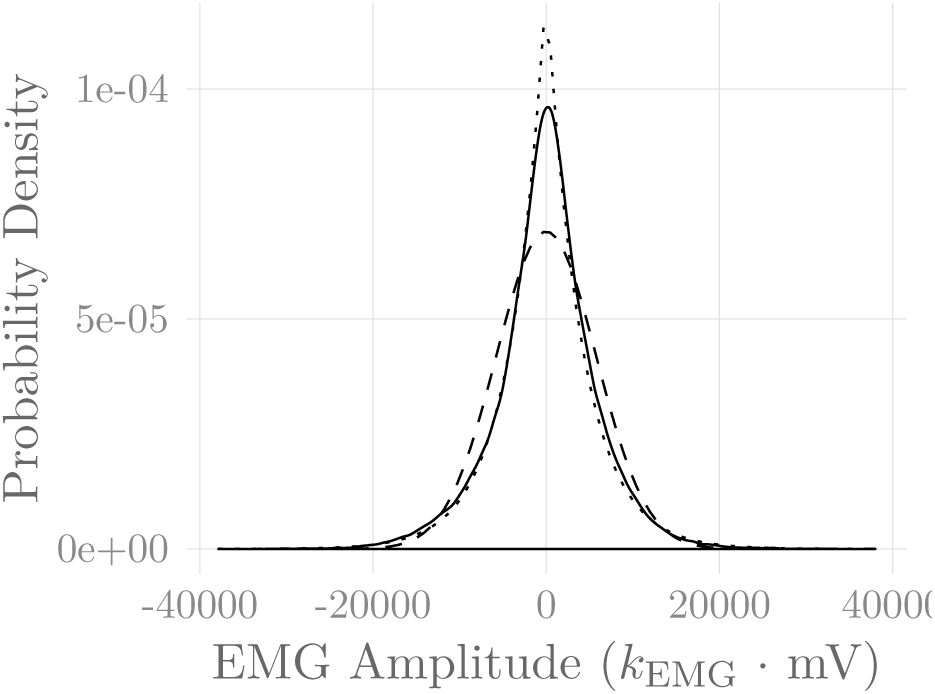
Amplitude distribution of the simulated sEMG signal over the course of a series of isometric contractions at progressively increasing activation levels (solid line). Superimposed are a Gaussian (dashed line) and a Laplacian (dotted line) distribution.

**Figure 6:**
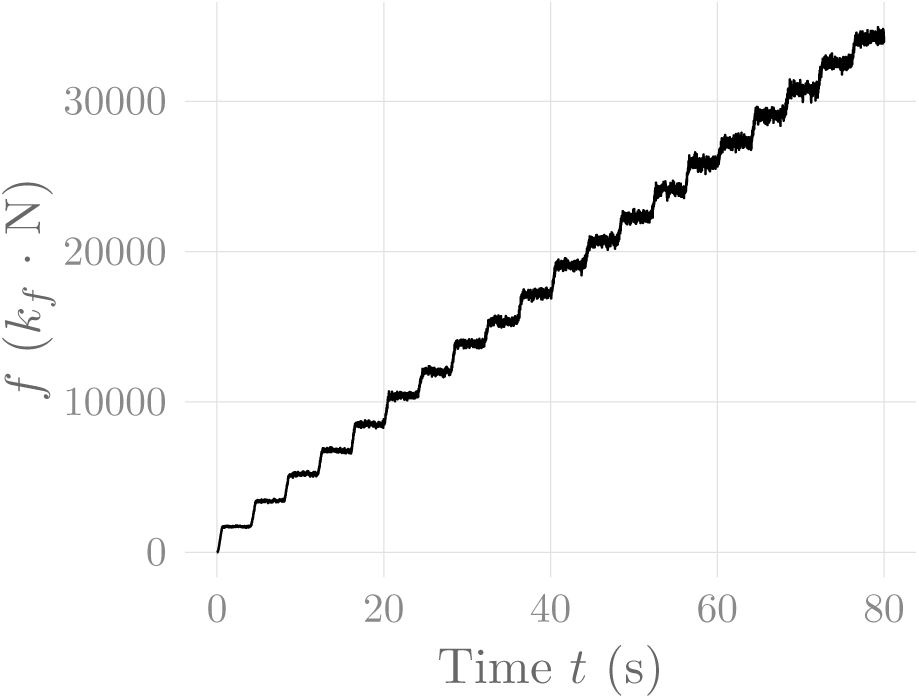
Simulated force signal of the simulated *recti* during a series of isometric contractions.

**Figure 7:**
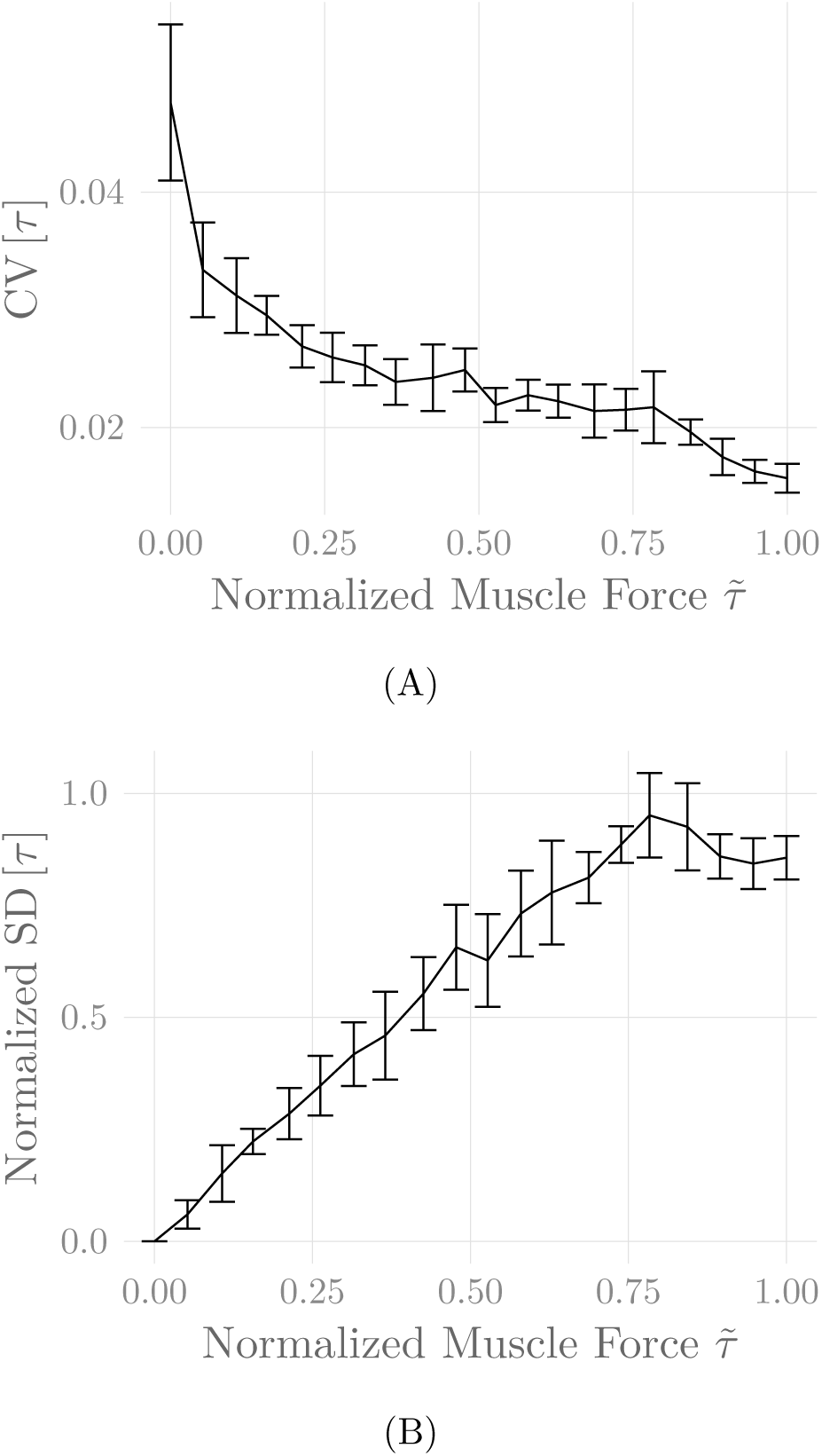
Coefficient of variation (A) and normalized standard deviation (B) of the simulated force signal of one belly of the simulated *recti* during a series of isometric contractions as a function of muscle activation. Shown are mean ± standard deviation of these values over seven simulation runs (including repeated MU placement).

**Figure 8:**
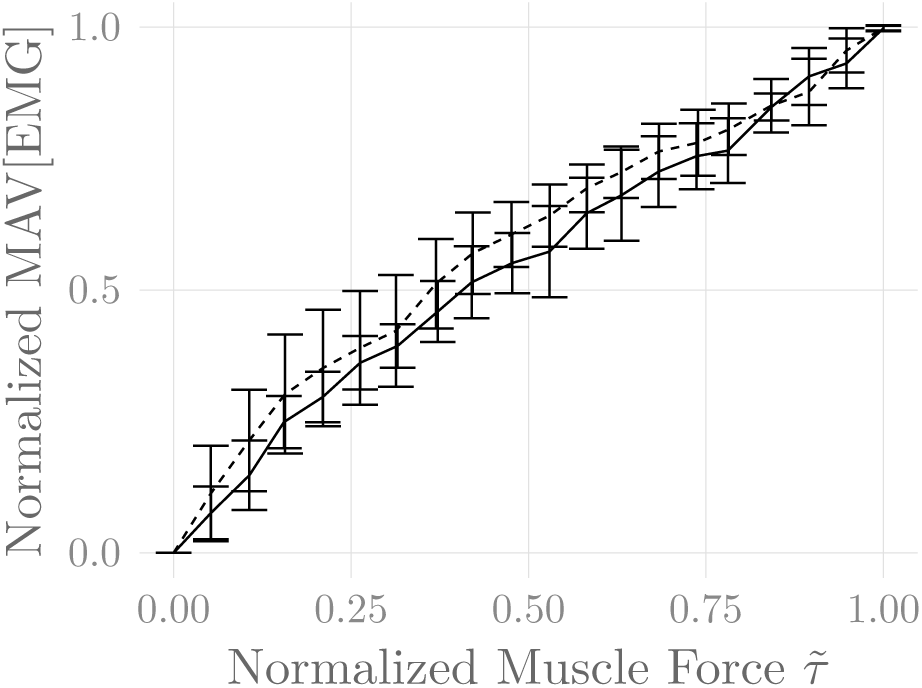
Steady-state sEMG-Force relation of the simulated *recti abdominis* over the course of the simulated constant-force isometric contractions. The first 250 ms of each contraction were discarded, and mean force and mean absolute value (MAV) of the EMG signal were calculated over the remainder of the contraction. Both values were ormalized to their minima and maxima over all contractions. Shown are the means of these values over five simulation runs (including repeated MU placement), both for the linear relationship described by equation (4) (solid line) and the square-root relationship described by equation (5) (dashed line).

## 4 Discussion and conclusion

In this article, a comprehensive mathematical model of surface EMG and force generation during voluntary isometric contractions in skeletal muscles has been described. The model consolidates and extends several previously proposed models for the different components of the physiological system that incorporate recent advances in understanding of physiology. In addition to combining previously isolated results in a unified model, a novel nonlinear input transformation is proposed which ensures that the generated muscle forces matches the target force level, and which alters the rate-coding and recruitment characteristics of the model in a physiologically meaningful way. Moreover, a more intuitive formulation of the EMG model of Farina and Merletti (17) is derived rigorously. Particular emphasis is placed on providing physiological justification for every component of the model.

The problem of input-output consistency of the simulated force signal has previously been solved by introducing a force feedback controller (7; 13; 54). While there certainly is a feedback component in neural force control, these controllers have not been designed to reproduce actual physiological force control. As long as it is not possible to reproduce the properties of the underlying physiological control system, it may thus be preferrable to use a static input nonlinearity for ensuring that the output force matches the desired force trajectory, instead of introducing further complexity into the simulation model in the form of a dynamic feedback loop. While the introduction of such a loop is inevitable once fatigueing contractions are simulated, the system may still benefit from the combination with a suitably-defined input nonlinearity, such as the one proposed in this article. This combination will be the subject of future research.

The proposed model may be useful, among others, for the simulation of realistic sEMG and force signals, for analyzing the sensitivity of the recorded signals to various physiological parameters, as well as to test signal processing algorithms on synthetic, yet realistic signals, measurements of which might be unavailable experimentally. Preliminary versions of this model have already been used successfully in previous publications of our group (2; 40) for the validation of algorithms for the separation of inspiratory and expiratory activity in sEMG measurements of the respiratory muscles. If desired, individual model components could be exchanged for other models of that particular physiological subsystem, e.g. to increase or reduce model complexity, or to take future physiological insights into account. Currently, effects due to the presence of muscle fatigue are not modelled; these could further be added to the model, e.g. by employing the metabolic model of Dideriksen et al. (13) and a force feedback loop (13; 54). For the simulation of limb muscles, the employed model of the volume conductor should be exchanged for the cylindrical model presented by Farina et al. (19). Finally, the restriction to isometric contractions could be resolved by implementing time-varying muscle geometry (and hence time-varying MU-electrode transmission paths), and by considering the force-length and force-velocity characteristics of skeletal muscles (56).

## Author contributions

Eike Petersen und Philipp Rostalski contributed to the conception of the work. The resulting mathematical model was derived and implemented by Eike Petersen under the supervision of Philipp Rostalski. Eike Petersen wrote the first draft of the manuscript. All authors contributed to manuscript revision and approved the submitted version.

## Acknowledgments

The authors would like to thank Marcus Eger, Drägerwerk AG&Co. KGaA, for many valuable discussions on the subject of this article.

## Data Availability

The simulated EMG, force, and impulse train signals of all six rectus bellies are available at the Dryad data repository (39) for one muscle with a linear EMG-force relationship and one muscle with a square-root relationship.

## Appendix A: Proofs

### Lemma 1.

*The expressions given in equations* (12) *and* (15) *to* (18) *are equivalent, assuming that ψ ∈ 𝒞∞*

*Proof.* Expanding all terms in equation (12) yields

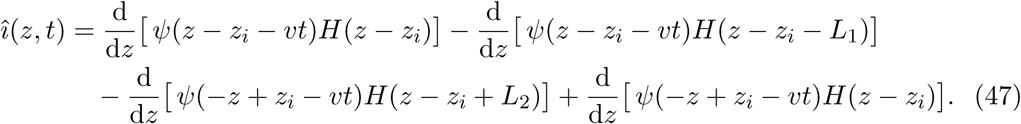

Note that the derivatives can only be understood in the sense of distributions, since the derivative of the Heaviside function *H*(*z*) can not be defined in the classical sense at *z* = 0. For some basic properties of distributions, refer to Appendix B.

Assuming *ψ* ∈ *C*^*∞*^ and recalling that ⟨(*ψH*)′, *ζ*⟩ = ⟨*ψ*′*H* + *ψδ, ζ*⟩ (refer to Appendix B), the duality ⟨*î*(*t*), *ζ*⟩ can hence be formulated as

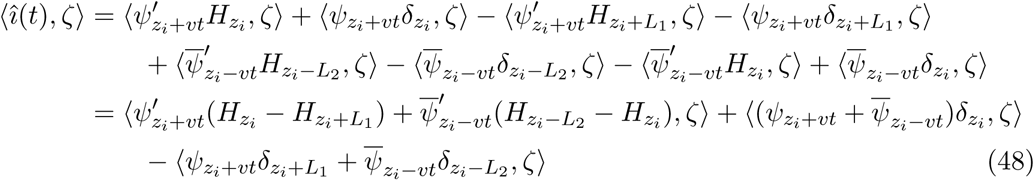

with notations 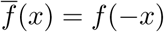and 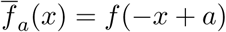 This is exactly equivalent to equations (15) to (18).

### Lemma 2.

*For compactly supported ψ*(*z*), *the IAP model given in equations* (15) *to* (18) *yields a formulation of î*(*z, t*) *that satisfies equation* (19).

*Proof.* For compactly supported *ψ*(*z*),

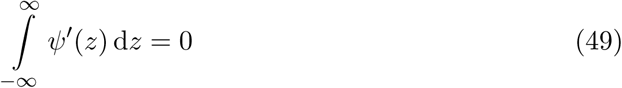

generally holds. Furthermore,

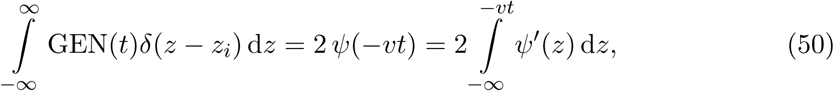

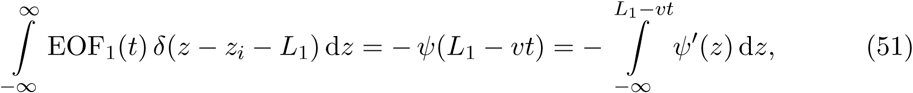

and equivalently for EOF_2_. Combining everything yields

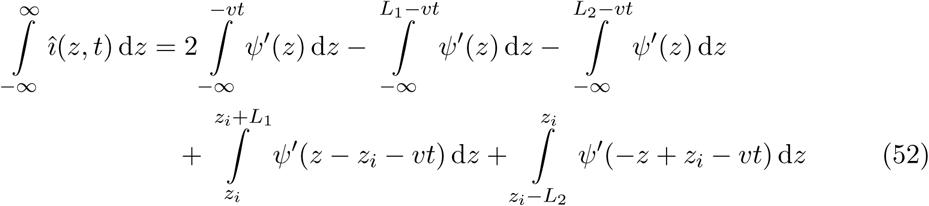

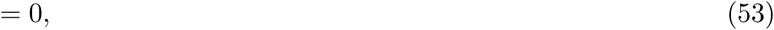

which concludes the proof.

## Appendix B: Distributions

A distribution *T* is a continuous linear mapping *T*: *D* → ℝ, where *D* is a given set of so-called test functions. In the following, the value of a distribution *T* acting on a test function *ζ* shall be denoted by the duality (⟨*T, ζ*⟩, with test functions chosen from the set *D* of compactly supported, smooth functions. Furthermore, two basic properties of distributions are

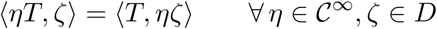

and

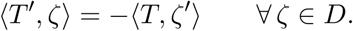

Moreover (following from the above), considering *η ∈ C*^*∞*,^

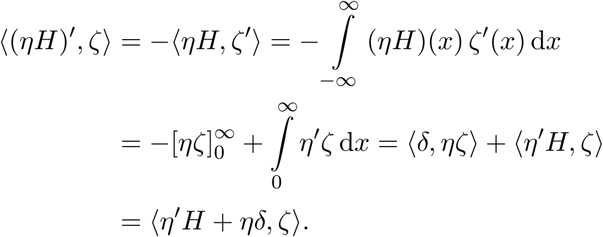

For more background on distributions (i.e., continuous linear functionals), refer to, e.g., Clarke (6).

## Footnotes

1 Note that – strictly speaking – the expression for *V*_*m*_(*z*) given in equation (14) does not result in a compactly supported *ψ*(*z*)*∈ C*^*∞*^, but equation (14) could trivially be modified to comply with this requirement, e.g. by convolution with a smooth and compactly supported mollifier. As this is a purely theoretical operation with no practical relevance at all, this has not been pursued here.

2 See previous footnote.

3 Note that in biological tissues, the charge carriors are ions, as opposed to electrons in electric wires (41, p. 25).

